# MOLECULAR LANDSCAPE OF OLD AGE MELANOMA BY SURVIVAL AND IMMUNOTHERAPY RESPONSE

**DOI:** 10.1101/2021.01.27.428444

**Authors:** Stephen P. Smith, Eduardo Nagore, Timothy Budden, Rajiv Kumar, Richard Marais, Caroline Gaudy-Marqueste, Amaya Virós

## Abstract

Melanoma mortality particularly affects older patients, and age is a powerful independent predictor of death. The pathogenic mutations and transcriptomic changes associated with poor survival in aged patients are not known.

We analyzed 5 cohorts of metastatic (N=324, N=18, N=66) and primary melanomas (N=103, N=30) to establish the effect of age on prognosis, identify age-specific driver genes and transcriptomic changes linked to survival and immunotherapy response.

We identify the pathogenic mutations and transcriptomic changes associated with poor survival by age, and show mutations in *BRAF, NRAS, CDKN2A* or *IDH1* identify metastatic and primary melanoma aged patients with worse outcome. In contrast, activation of immune-regulatory pathways is a hallmark of long-term survival. We tested if mutations in genes linked to poor outcome are associated to immunotherapy responders, exploring combinations of agespecific mutations in metastatic immune checkpoint inhibitor aged responders. Strikingly, aged patients with *BRAF, NRAS, CDKN2A* or *IDH1* mutations and high tumor mutation burden treated with immunotherapy have an improved median survival of 12 months. These data highlight the molecular landscape of melanoma varies by age, and age stratification can refine prognosis and therapy rationales. A set of mutations identifies patients at highest risk of death who are likely immunotherapy responders.

## Introduction

The incidence and mortality of melanoma increases with age (Balch 2015; Green and Olsen 2017; Hillen et al. 2018) and >80% of melanoma deaths affect patients older than 59 (Balch 2015; Tsai et al. 2010). Older patients more frequently present with thicker primary tumors, and additional characteristics of poor prognosis: ulceration, elevated mitotic rate, and early visceral metastasis (Balch 2015). However, even after taking the main prognostic factors into account, there is a survival discrepancy between the elderly and young (Balch 2015; Cavanaugh-Hussey et al. 2015), and age is the most important independent marker of adverse outcome together with tumor thickness (Balch et al. 2013; Balch et al. 2001). The large prevalence and mortality of disease affecting the aged population is not reflected in current trials, and therefore identifying markers of prognosis and therapy response to stratify aged patients, at higher risk of death, is paramount.

Older patients clearly stand to benefit from immunotherapy, and some reports suggest aged patients may present better response rates to immune checkpoint inhibitors (ICI) than younger patients (Joshi et al. 2018; Kugel et al. 2018; Perier-Muzet et al. 2018; Ribas and Wolchok 2018; De Rosa et al. 2018; Schadendorf et al. 2018). ICI are currently being tested in the adjuvant, early stages of disease (Eggermont et al. 2018b; Eggermont et al. 2018a), but there are no validated rationales to stratify patients based on molecular evidence of risk of progression or response to therapy.

Individual mutations and transcriptomic changes are poor predictors of survival and therapy response (Akbani et al. 2015; Berger et al. 2012; Chen et al. 2016; Hodis et al. 2012; Roh et al. 2017; Weinstein et al. 2014), but the impact of age-specific mutations on survival and treatment has not been studied systematically previously. We hypothesized that poor outcome of older patients with melanoma can be predicted from specific molecular changes, and that age-specific molecular changes can be used to stratify patients for immunotherapy. In this study we investigated the association between age, age-specific molecular changes and survival in primary and metastatic melanoma. We tested if molecular markers linked to poor outcome and age can identify patients who respond to ICI.

## Results

### Old age is a primary determinant of outcome in malignant melanoma

We first explored the relationship between age and survival analysing data from the Cancer Genome Atlas SKCM (cutaneous melanoma). For an initial assessment, we compared metastatic melanoma patients by an age cutoff of >59 years old and <=55 years old, excluding intermediate age groups (Figure 1A, Table 1A). In this metastatic population (n=324, median follow-up 47.5 months, 137 patients deceased) age >59 was significantly associated with poorer overall survival compared with age <=55 (p<0.0001, median survival 62 vs 144 months). Multivariate stepwise regression (Table 1B) showed that age was the most significant determinant of outcome (p=0.001) followed by stage at diagnosis (p=0.025). To confirming that age specifically affects the course of melanoma we investigated the impact of age on progression free survival (PFS, Figure 1B) and found a similar effect (p=0.013, 48 vs 143 months median survival). To confirm the relationship between age and survival is not limited to a single age cutoff, we confirmed our survival results using all age cutoff for ages 50-70, using the entire population, obtaining similar significance.

**Figure 1.**
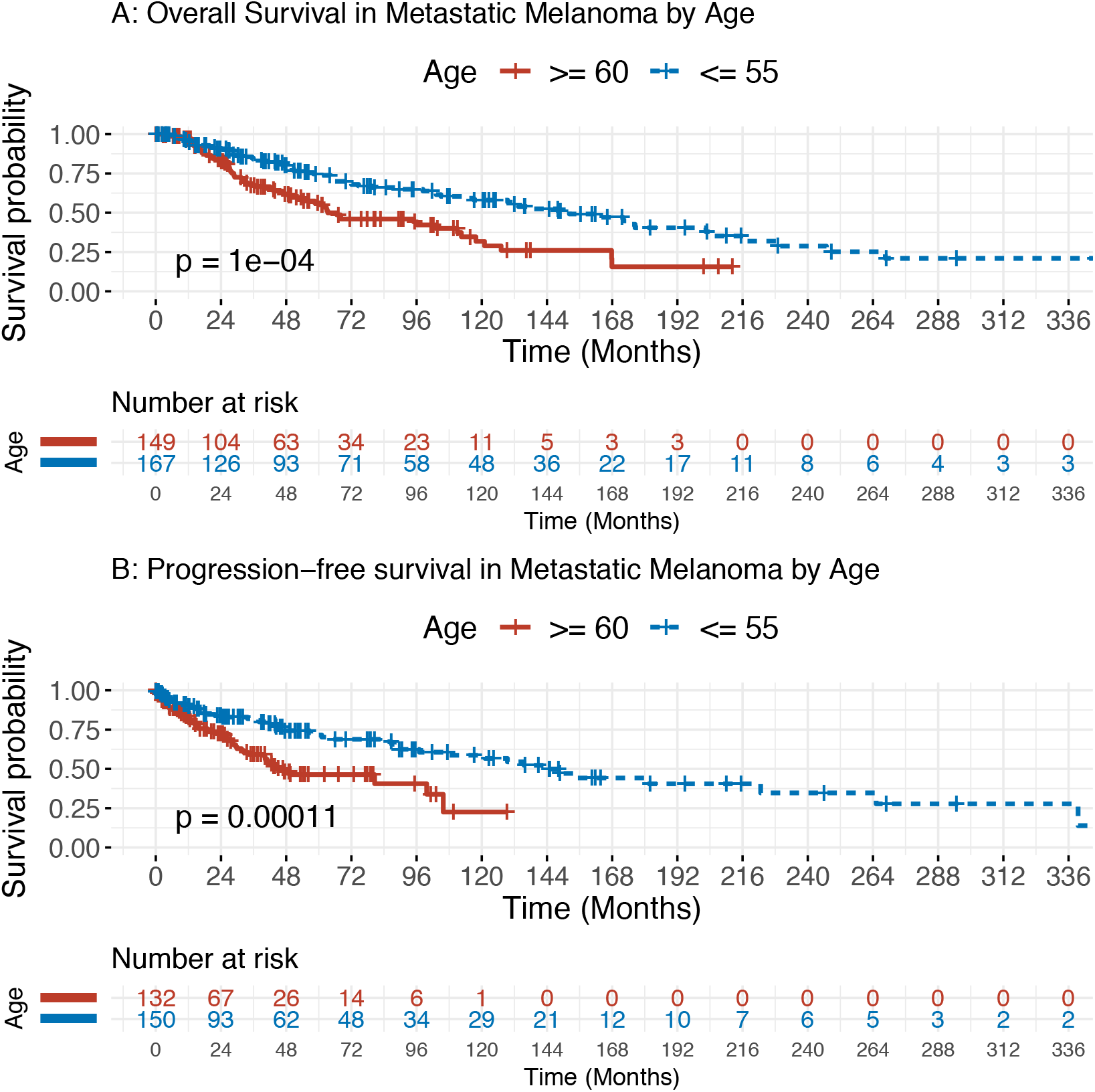
Overall and progression-free survival of patients with metastatic melanoma from TCGA, stratified by age. All patients with metastatic melanoma in the TCGA SKCM cohort were included and their survival probability calculated. Log-rank testing established a significant overall **(A)** and progression free **(B)** survival advantage in the group of age <55.

**Table 1.**
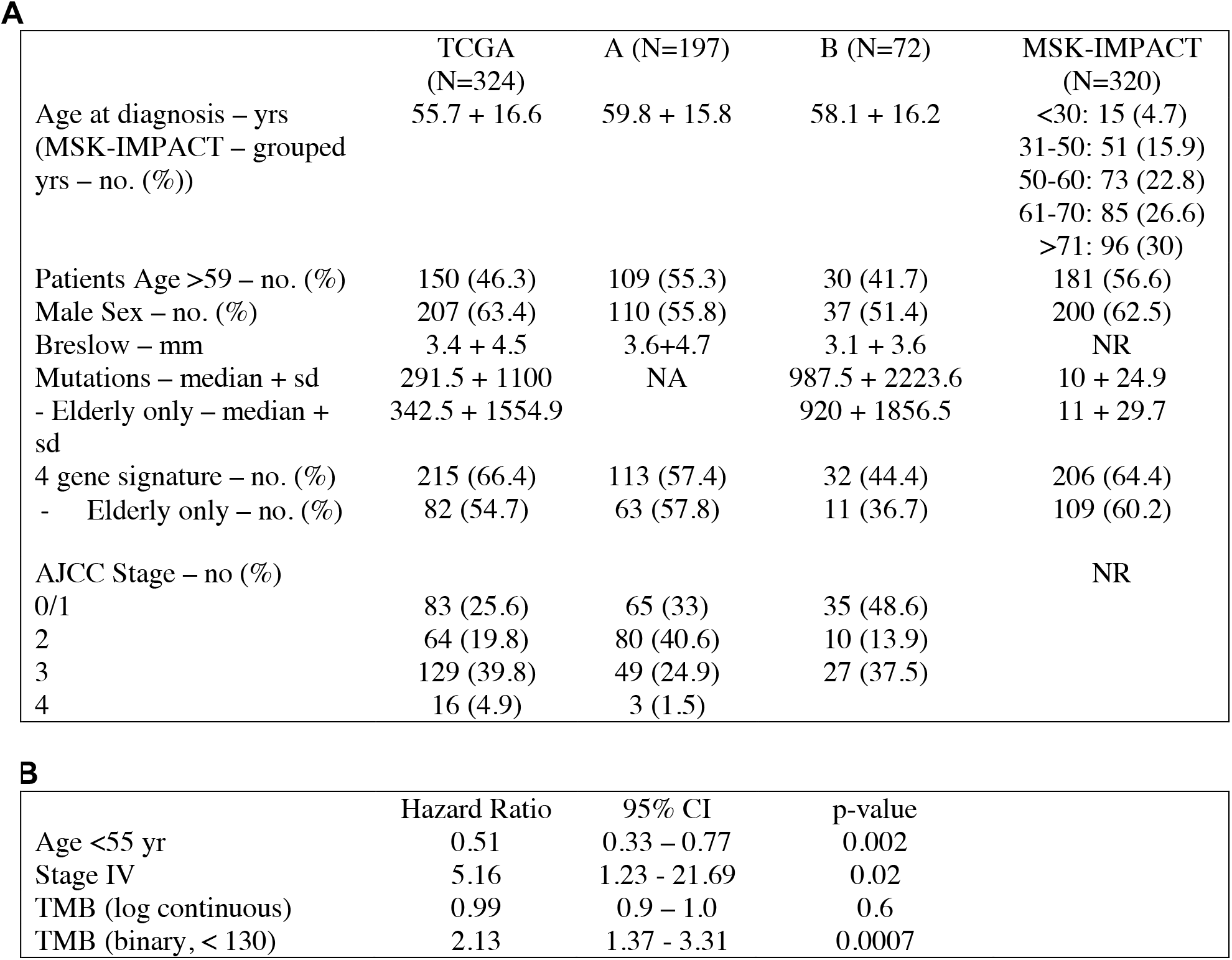
(A) Demographic, clinical and molecular characteristics of patients in the 4 tested cohorts, and (B) Results of multivariate regression analysis of TCGA patient survival. Age, Breslow = mean + sd. Exact ages and staging information not given in MSK-IMPACT. Breslow thickness and stage not recorded in MSK-IMPACT. Mutation count: all mutations in TCGA, non-synonymous variants from selected gene panel in MSK-IMPACT. NA: not available.

### Genomic markers of poor outcome in metastatic melanoma of older patients

Older patients carry a significantly higher tumour mutation burden (TMB) than younger patients (median 335 old, 274 young, p<0.0001), and previous work reports a survival benefit in melanoma patients with high TMB(Gupta et al. 2015; Trucco et al. 2018). However, we found TMB was not significantly associated with survival after multivariate regression, or when analyzed as a continuous variable in multivariate regression (logarithmic or unchanged, Table 1B, Supplementary methods). Only when TMB was considered as a binary variable (high or low defined by a cut-off of 130 mutations), did we find an association with improved survival (Gupta et al. 2015). This finding remained true for PFS and for all age cutoffs in the range of 50-70.

We explored the distinct molecular characteristics of older age melanoma (>59). Mutational signature decomposition demonstrated that the mutations in all ages were overwhelmingly consistent with ultraviolet radiation (UVR)-mediated DNA damage (Supplementary Figure 1A, 1B) as expected. We identified areas of hypermutation, where multiple consecutive mutations occur in short stretches of the genome, more frequently in older patients (Figure 2A), indicating that elderly patients accumulate more mutations overall and in specific genomic regions that are more susceptible to UVR-induced damage. Furthermore, the genes affected by mutations in these regions were more numerous and varied than in younger patients (Supplementary Figure 2).

**Figure 2.**
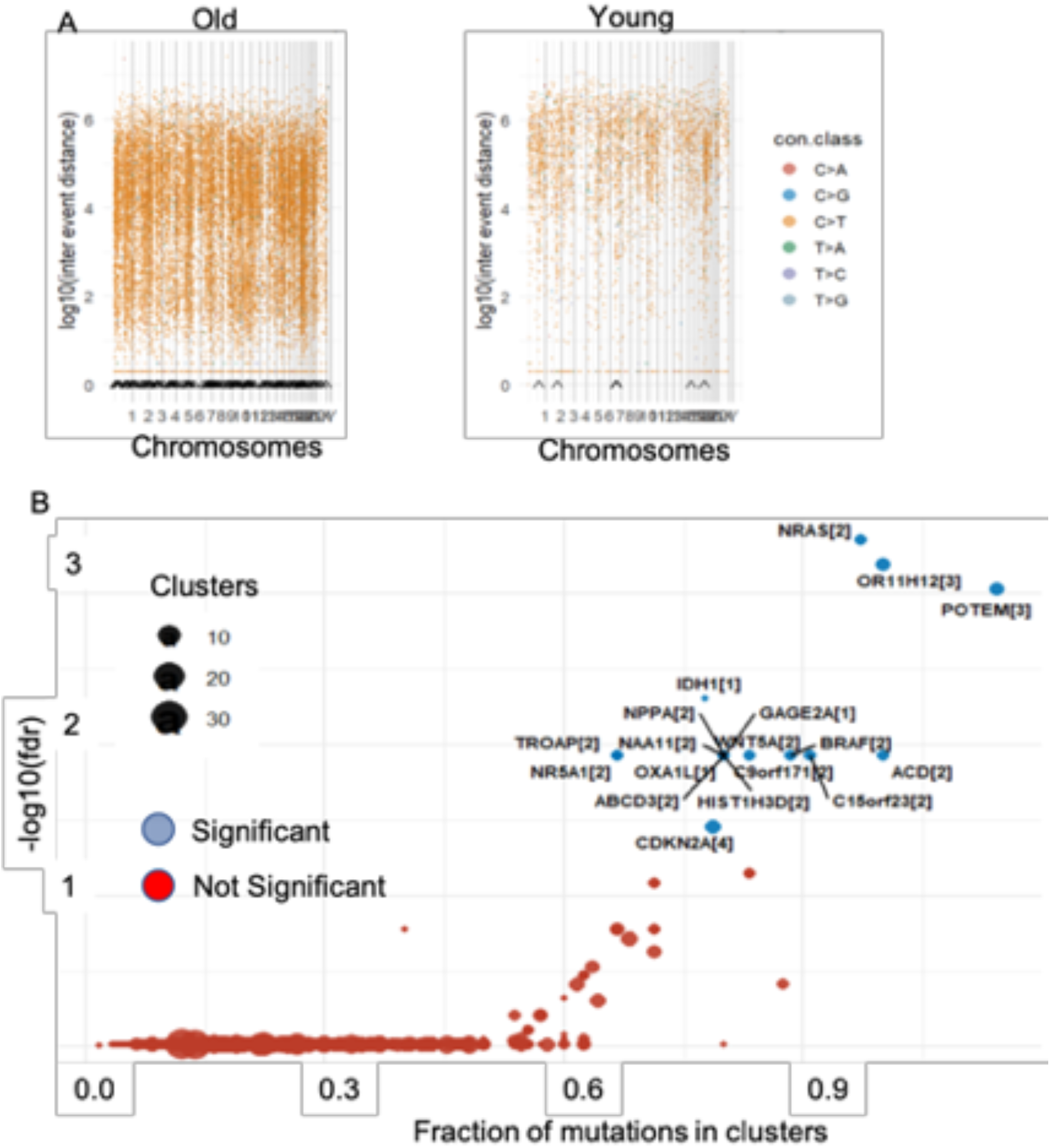
Mutational characterisation of metastatic melanoma (TCGA). (**A**) Top 2 panels show rainfall plots of total mutations across the genome (x) by the distance from the previous mutation (y) illustrating the greater density of mutations in the elderly (left panel). Black arrows on the x axis indicate points of hypermutation. (**B**) Results of the oncodriveCLUST algorithm, which identifies 18 driver genes in elderly melanoma, brackets indicate the cluster of mutation for that gene.

To identify the mutational drivers associated with older age melanoma, we used the oncodriveCLUST algorithm (Tamborero et al. 2013) which accounts for background mutation density and gene length to identify key “driver” mutations (Figure 2B). We identified 18 driver genes in aged samples and 11 drivers in young patients (Supplementary Figure 3), and only BRAF and NRAS were found in both groups. Surprisingly, mutations in any of the 18 putative driver genes of older melanoma were significantly associated with poor prognosis (p= 0.013, Supplementary Figure 4).

We next studied the transcriptional profile of old age cutaneous melanomas by survival. For this, we first compared the differentially enriched pathways in long-term survivors (≥2000 days) to short-term survivors (<2000 days) in metastatic melanomas, and then performed unbiased gene set enrichment analysis (GSEA (Subramanian et al. 2005)). A total of 870 genes were significantly differentially expressed (FDR *q*<0.1) between groups, with 297 genes expressed higher in long-term survivors and 573 genes higher in short-term survivors. Remarkably, genes expressed in long-term survivors were enriched with genes from many immune related pathways involved in cancer control and immunotherapy response (Figure 3A). We found activation of interferons and cytokine signaling pathways, and PD-1 signaling genes, including eight HLA gene and the antigen presentation genes TAP1, and B2M. Additionally, long-term survivors had a significantly higher CD8 T cell score than the short-term survivors (p = 0.048, Figure 3B)).

**Figure 3.**
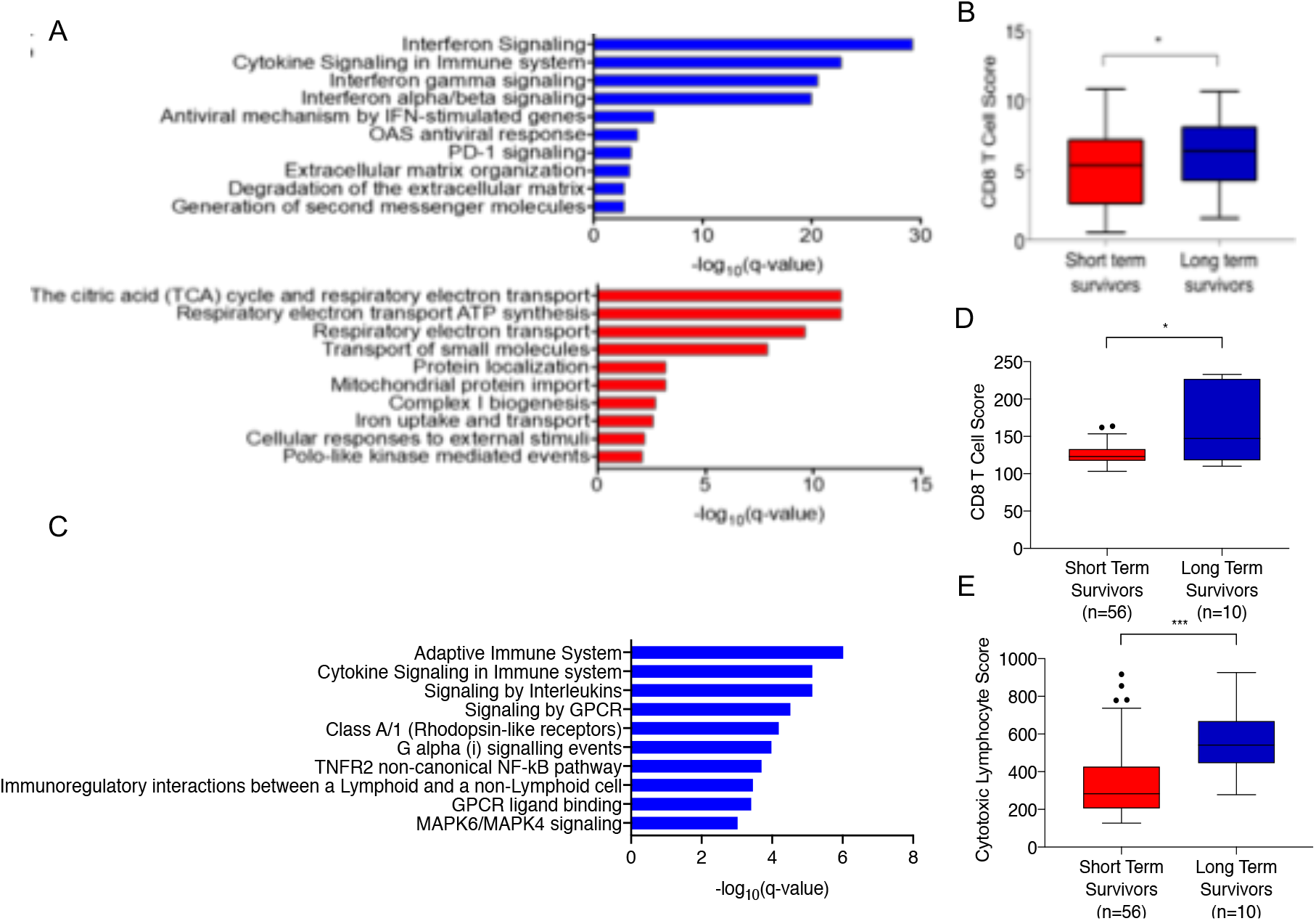
(**A**) Cell signaling pathways enriched in long-term (blue) or short-term (red) survivors with metastatic melanoma from the TCGA cohort. **(B)** CD8 T-cell score derived from TCGA bulk RNA sequencing in metastatic melanoma showing a significant (p < 0.05) difference between long and short-term survivors in this cohort. (**C**) Cell signaling pathways enriched in long-term (blue) survivors with metastatic melanoma from the Lund cohort. **(D)** CD8 T-cell score and **(E)** cytotoxic lymphocyte score derived from Lund microarray data in metastatic melanoma patients showing a significant (p < 0.05, <0.001) difference between long and shortterm survivors in this cohort.

Short-term survival genes were enriched for several metabolism pathways, enriched for the electron transport chain and oxidative phosphorylation pathways, with genes encoding the subunits for cytochrome c oxidase, NADH:ubiquinone oxidoreductase, and ubiquinol-cytochrome c reductase complexes. Transport of small molecules pathway was also significantly enriched with 21 solute carrier family genes (Figure 3A).

We next validated the transcriptional profile associated to improved survival in old age in microarray data of an independent cohort of 66 metastatic melanoma samples of patients >59(Cirenajwis et al. 2017). As before, we compared the differentially enriched pathways in long-term survivors (≥2000 days) to short-term survivors (<2000 days) and confirmed genes expressed in long-term survivors were enriched for immune related pathways involved in cancer control and immunotherapy response (Figure 3C). Furthermore, we confirmed longterm survivors had higher CD8 T cell scores and cytotoxic lymphocyte score than the shortterm survivors (p = 0.03, p = 0.0001; Fig 3D, 3E).

### Genomic markers can be used to predict survival in primary melanoma

A key clinical need is to stratify patients for risk of death as early as possible in their clinical journey. The widespread increase and accessibility to DNA sequencing in routine pathology services for multiple cancers prompted us to test if any of the predicted oncogenes we discovered linked to poor outcome in metastatic melanomas of the TCGA could be easily used in primary melanoma as a biomarker for older patient outcome. We selected a core set of four well-characterized, key melanoma oncogenes (BRAF, NRAS, IDH1 and CDKN2A) from the 18 driver genes associated to poor outcome in the aged metastatic cohort because they are routinely sequenced in current cancer sequencing panels. The TCGA primary melanoma cohort includes primarily highly aggressive, thick, ulcerated primary tumors; with short follow-up (Supplementary Table S1), so we tested if mutations in BRAF, NRAS, IDH1 and CDKN2A can predict outcome of aged primary melanoma patients in two separate and independent cohorts (Table 1A). We confirmed that DNA mutations in one or more of the 4 genes (BRAF, NRAS, IDH1 and CDKN2A) significantly and powerfully predicted poor disease-specific outcomes in patients aged >59 with primary melanoma (Figure 4), and confirmed the predictive power of mutations in these 4 genes in the other age cutoffs remains significant. Importantly, the mutation frequency of these 4 genes rises with age, so it is in the older population where it has the strongest predictive power. In contrast, a set of alternative genes that can more accurately predict outcome in younger patients (Supplementary Figure 4).

**Figure 4.**
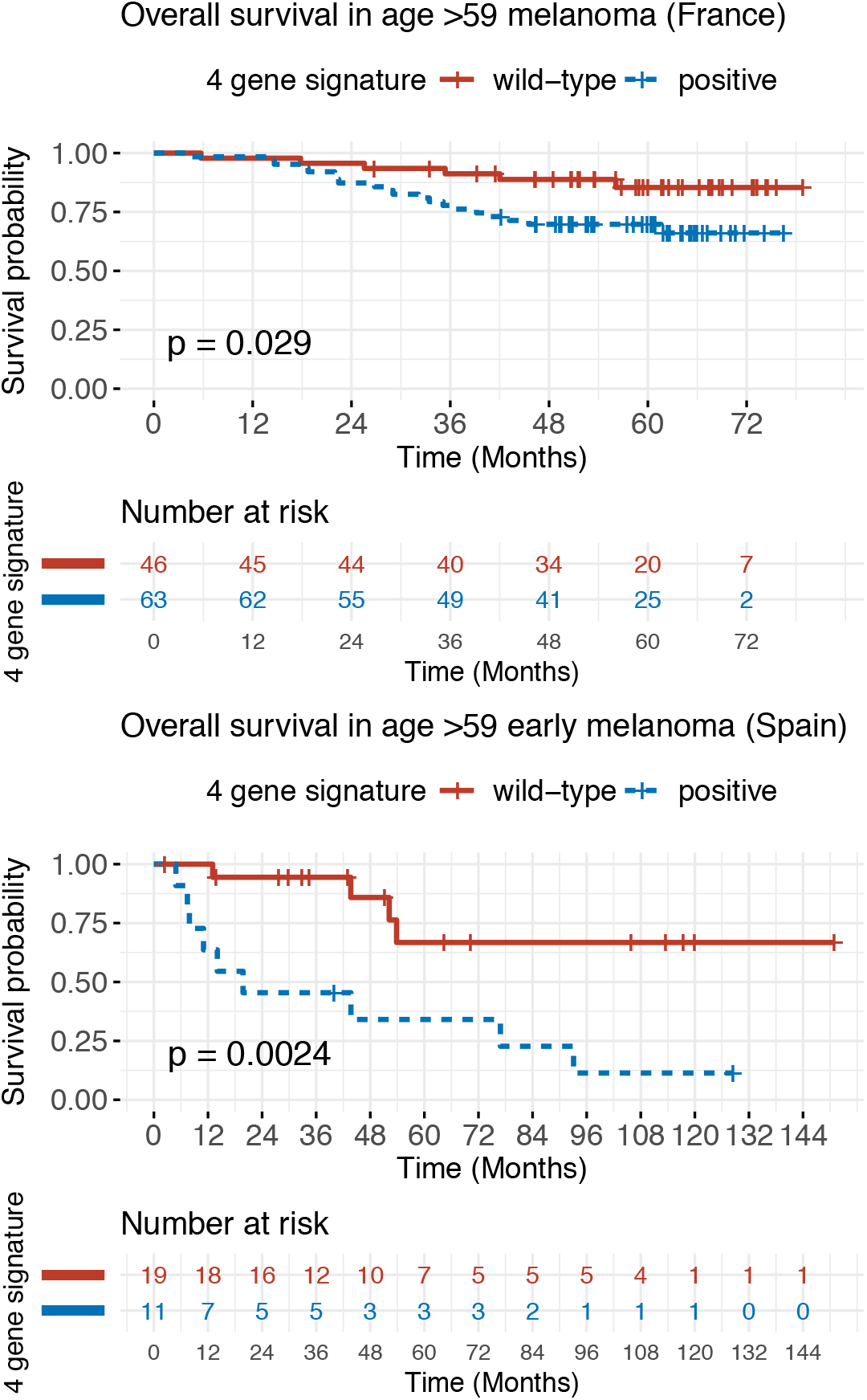
Overall survival in melanoma patients of age >59 from independent cohorts (Top: A cohort (France), Bottom: B cohort (Spain)) with primary melanomas. Both panels show the significant survival difference between patients with or without mutations in the 4-gene core driver set identified from TCGA data. Despite the different baseline characteristics of the cohorts the gene mutation signature significantly predicts poor outcome in both cohorts.

### Genetic mutations and tumor mutation burden in aged patients identifies responders to immunotherapy

Prediction of response to ICI for high-risk, early and late stage melanoma is a pressing clinical requirement and to date there are no reliable pre-treatment genomic biomarkers to select patients at highest risk of death and a higher likelihood of response. Recent evidence shows a higher TMB leads to a greater likelihood of neo-antigen-driven response in the melanoma population not selected for age (Goodman et al. 2017; Johnson et al. 2016a; Liu et al. 2019; McGranahan et al. 2016; Rizvi et al. 2015; Snyder et al. 2014); and additionally, the presence of two of our core signature genes NRAS (Johnson et al. 2015) and CDKN2A mutations (Helgadottir et al. 2018) in metastatic melanoma identify subgroups of patients with improved rates of immunotherapy response. Previous studies have shown a link between TMB and immunotherapy response (Gibney et al. 2016; Johnson et al. 2016b; Liu et al. 2019; Roszik et al. 2016), however data from the large-scale MSK-IMPACT study has demonstrated that although TMB is a predictor of survival in immunotherapy-treated patients across all cancer types, it is not a powerful predictor of immunotherapy response in melanoma (Samstein et al.). We reasoned that although genetic damage alone is a limited approach to stratify patients, age and TMB are significantly correlated (Supplementary Figure 5), and combining genetic markers linked to outcome by age with TMB could improve our current approach to stratification. To test this hypothesis, we investigated the relationship between age, TMB, response to immunotherapy and the presence of mutations in BRAF, NRAS, IDH1 or CDKN2A, which strongly predict survival.

We found that independently, the mutations in the four-genes and the TMB did not predict response; but when combined, powerfully identified responders to ICI in the older population (Figure 5). The median survival difference between older, mutation-positive and mutation-negative patients with high TMB was approximately 12 months, and the difference between mutation-positive, high TMB and low TMB patients was 10 months (Figure 5A, 5B, 5C). To test this association is specific to the older population, we tested our approach in young patients, and found neither TMB nor mutations alone or in combination identified immunotherapy responders (Figure 5D). Finally, we found no differences in expression of the previously described interferon pathway expression signature of ICI response (Ayers et al. 2017) between patients with mutations in the 4 genes and mutation-negative patients (p > 0.4), or with age (r2 = 0.03, p > 0.6).

**Figure 5.**
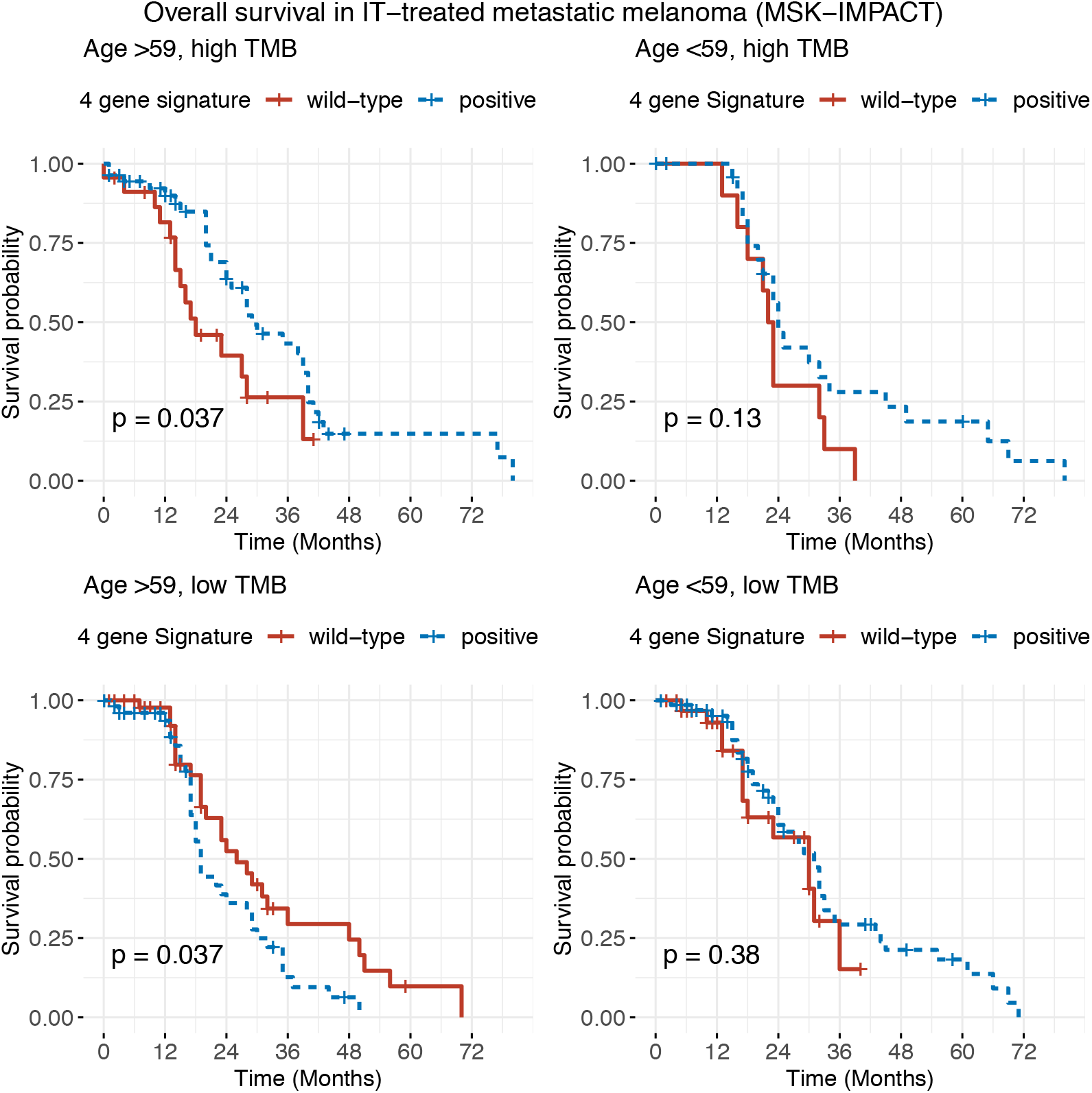
Overall survival in immunotherapy treated metastatic melanoma from the MSK-IMPACT study stratified by age and the 4 gene driver signature. Data from the MSK-IMPACT study were re-analysed to calculate the survival probabilities in old or young patients with or without the 4 gene driver mutation signature who show TMB higher or lower than median level. The data show that in old (but not young) patients a high TMB and the presence of the 4 gene driver signature is a positive prognostic marker for survival with ICI therapy, but with a low TMB the prognosis is worse. In both groups the difference is clinically (and statistically) significant.

## Discussion

The clinical and health economic need for simple, reliable tests to identify patients with melanoma whose trajectory is likely to be worse, and those who are most likely to respond to ICI therapies is clear. Untreated older melanoma patients have a significantly worse outcome, and the vast genomic differences that we show here following age-specific dissection of molecular data, supports the view that melanoma in elderly patients comprises a distinct group. Older patients respond to ICI (Joshi et al. 2018; Kugel et al. 2018; Perier-Muzet et al. 2018; De Rosa et al. 2018), and importantly, there are tumour immune-state response predictors (Balachandran et al. 2017; Chowell et al. 2018; McGranahan et al. 2016; Rooney et al. 2015; Topalian et al. 2012) of poor outcome that are more prevalent in the young (Kugel et al. 2018). High levels of accrued DNA damage, an established biomarker of ICI response in other cancer types(Samstein et al.), is highly present in aged melanomas.

Our work defines the molecular changes that identify aged patients at highest risk of death. We show there is an age-specific mutational landscape of oncogenic drivers that predict a worse outcome in both early and late-stage melanomas. Moreover, we describe the transcriptomic features that exist in untreated, metastatic samples that are linked to survival, finding a highly significant up-regulation of immune related pathways and CD8 T cells in long term aged survivors. Intriguingly, previous studies investigating the transcriptome in ICI metastatic melanoma responders prior to therapy identify the upregulation of immune pathways, in common with our study, as predictive of improved outcome. Both studies show the immune features of the tumour are critical for disease control, either to prolong survival without therapy, or to enable response to ICI. Finally, we show a simple set of mutated genes in aged metastatic melanomas, combined with TMB, discerns patients with a 12-month median longer response to ICI.

This study is limited by being retrospective in nature but strengthened by using prospectively collected “real-world” data. The five cohorts investigated differ in the proportion of patients at different stage of disease and in the methodology used to identify mutations and transcriptomic changes. This makes comparison between them challenging, but despite these limitations the robustness of the findings is encouraging – as is the fact that the data required to test the applicability of the 4 mutated genes were available from patients seen in routine clinical practice, in separate countries.

Differentiating between groups based on a continuous variable such as age is challenging, and we chose to use an arbitrary pair of limits to define “old” and “young” patients. We have explored varying these cutoffs across a wide range, as well as including all patients. The results remain consistent with those changes, and further prospective studies should address the most useful, robust cutoff considering disease prevalence by age and mutation prevalence in larger cohorts. Additional studies could also address the specific mutations that confer the most risk, to develop a simple set of predictive markers that can be applied routinely in clinic.

The genetic predictors of decreased survival in old age melanoma additionally identify the subset of metastatic patients who will derive the most benefit from checkpoint inhibitors. Mutations in the 4 genes (BRAF, NRAS, IDH1 or CDKN2A) represent a robust, simple set of analyses that can be easily incorporated into clinical practice to identify aged patients at genetic high risk of melanoma progression, and when combined with TMB, can identify the patients who are most likely to respond to immunotherapy. Given the cost and toxicities of ICI, this study provides a biomarker that can stratify patients, predict response and support therapy decisions. This study addresses an area of high unmet clinical need in a group of patients who stand to derive significant benefit from its findings.

## Methods

### Cohorts

The exploratory analysis of age and genetic changes was undertaken in the TCGA dataset for metastatic patients age >59 vs <=55(Akbani et al. 2015; Weinstein et al. 2014) and (https://www.cancer.gov/about-nci/organization/ccg/research/structural-genomics/tcga). Primary melanomas >59yo were from A cohort (n=109 patients) and B cohort (n=30 patients, Table 1, Supplementary Table S1). The effect of mutations and tumor mutation burden in immunotherapy-treated patients was investigated in the MSK-IMPACT study (n=181 patients >59yo (Samstein et al.)).

The experimental age cut-offs age >59 or <=55 were decided *a priori* to obtain more clear-cut young and old age categories. To ensure this did not introduce bias we then analysed the data using a simple <59 vs >60 cut-off and a wide interval of <50 vs >70. To validate the findings we studied clinical and sequencing data collected in cohort A: La Timone University Hospital, (Aix Marseillle, France) and cohort B: Instituto Valenciano de Oncología (Valencia, Spain) (Gaudy-Marqueste et al. 2014). Relevant institutional review boards and ethics committees at both institutions in both countries approved molecular research and all human participants gave written informed consent. The primary cohorts are prospectively collected and unselected (supplemental table S1), subjected to targeted deep sequencing at a depth of x100 of 40 melanoma genes. The sequencing panel includes genes mutated in >10% of TCGA samples or mutated in less than <10% of samples but known cancer drivers. The 4-gene signature is significantly less frequent in the primary cohorts (p= 0.0008 B cohort, p=0.048 A cohort) than in the metastatic melanoma TCGA cohort, as you would expect for unselected populations of primary melanoma that, in contrast to metastatic TCGA samples, are not enriched for samples with poor prognosis. However, the proportions are not significantly different between any of the primary cohorts (p>0.1 in all comparisons).

### Statistics

Regression modelling of survival in the metastatic melanoma (TCGA) cohort was conducted using Cox proportional hazards modelling in R implemented in the Survival package. We included age, sex, patient weight, Breslow tumour thickness of original primary tumour and somatic variant count. To derive the minimum model, we excluded the least statistically significant variable at each step until all variables were significant (p<0.05) (supplemental table S2). To explore the effects of variant count in the model we ran the above sequence initially with variants as a log continuous variable, and then as a binary (high/low) variable with a series of cut-off values (25, 50, 100, 110, 150, 150, 500). Of these only 100 and 110 showed a significant impact in the final model and both were less significant than age in those cases.

We analysed the mutation data using the oncodriveCLUST algorithm (Tamborero et al. 2013) as implemented in the maftools package in R to discover key driver mutations. OncodriveCLUST considers the background mutation rate, gene length and the clustering characteristics of mutations identified in order to derive a confidence value for labelling a gene as a driver as opposed to a passenger.

### Gene expression

For the TCGA data, differential gene expression analysis was performed using the DESeq2 (version .28.1) (Love et al. 2014) package in R (version 4.0.0, RStudio v1.2.5003, RStudio Inc). RNA-seq count data for the TCGA SKCM cohort was acquired using the TCGABiolinks package (version 2.16.0 (Colaprico et al. 2015)). The count dataset was filtered to remove genes with less than 1000 counts across all samples. Differential expression analysis was run between long and short-term survivors in the aged (>59 years) metastatic cohort. Long-term survivors were defined as patients with overall survival time 2000 days or greater, compared to patients who died <2000 days. For pathway enrichment analysis genes that were significantly differentially expressed (FDR p-value < 0.1) were compared against the Reactome Database (Jassal et al. 2020) using the Molecular Signatures Database v7.0 (Msigdb, Broad Institute, https://www.gsea-msigdb.org/gsea/msigdb/index.jsp).

The Lund expression data (Gene Expression Omnibus GSE65904) of 214 melanomas(Cirenajwis et al. 2017), includes 66 metastatic patients with long term survival data. We tested the differential expression between aged long and short-term survivors (Limma package (version 3.44.1) in R). MCPCounter package generated cell population scores (Ritchie et al. 2015). Cell population scores were generated using the MCPCounter R package (version 1.2.0 (Becht et al. 2016)) in the TCGA SKCM RSEM normalised RNA-seq expression data. CD8 T cell scores were compared between long and short-term aged metastases using Mann-Whitney in GraphPad Prism (version 7.01).

### Response to Immunotherapy

Data from MSK-IMPACT (Samstein et al.) was downloaded from cBioPortal (http://www.cbioportal.org/study/summary?id=msk_impact_2017) and reanalyzed (Zehir et al. 2017). To determine high vs low TMB we investigated the centile groupings within the melanoma group and used the third decile (i.e. those in the highest 30% TMB) as high. As did the original MSK-IMPACT, we tested all deciles (Samstein et al.). 30% was the inflection point at which the response curve separation began to accelerate. Any cutoff from 30% to 10% results in significance (with increasing magnitude as the cutoff becomes more stringent) – we chose a cutoff at 30% to include as many patients as possible.

### Data availability

Anonymized data are available without restriction. The analytical code is available from the authors without restriction.

## Abbreviations

(ICI): Immune checkpoint inhibitors
(PFS): Progression free survival
(TMB): Tumour mutation burden
(UVR): Ultraviolet radiation
(GSEA): Gene set enrichment analysis
(TCGA): The Cancer Genome Atlas

## Acknowledgments

AV is a Wellcome Beit Fellow and personally funded by a Wellcome Trust Intermediate Fellowship (110078/Z/15/Z), Cancer Research UK (A27412), Leo Pharma Gold Award and Royal Society (RGS\R1\201222). SPS is funded by the Medical Research Council UK (MR/R001146/1). EN is funded by Fondo de Investigación en Salud (FIS) PI15/01860, Instituto Carlos III, Spain. RK is funded by TRANSCAN (01KT15511), German Ministry of Education and Research (BMBF) and DKFZ. RM is funded by Cancer Research UK (A27412) and Wellcome Trust (100282/Z/12/Z). CG-M is funded by the French Dermatology Society, Collège des Enseignants en Dermatologie de France (CEDEF) and UNICANCER France. We acknowledge the generous contribution of APHM Biobank (France) and IVO Biobank (Spain). Bioresources were provided by the Biological Resources Centre of the Assistance Publique Hôpitaux de Marseille, (CRB AP-HM, certified NF S96-900 & ISO 9001 v2015), from the CRB-TBM component (BB-0033-00097), and from the Biobank of the Instituto Valenciano de Oncología, Valencia, Spain.

## Supplementary Material

### Supplementary Methods

**Figure S1.** Mutational signature decomposition old and young melanoma samples from TCGA.

**Figure S2.** Summary of the types and frequencies of mutations in metastatic melanoma samples from old or young patients in TCGA.

**Figure S3.** OncodriveCLUST analysis of metastatic melanoma in age <55.

**Figure S4.** Overall survival in metastatic melanoma in TCGA.

**Figure S5.** Correlation between age and TMB across in the TCGA and MSK-IMPACT cohorts.

**Table S1.** Full clinical details of the patients included in the analysis of the elderly (age >= 60) A and B primary melanoma cohorts.

**Table S2.** Multivariate regression using a cox proportional hazards model.

### Supplementary Methods

The exploratory analysis of the effect of age and mutations on survival were undertaken in the TCGA dataset for metastatic patients age >59 vs <=55. The cohorts of primary melanomas >59yo were from A cohort (n=109 patients) and B cohort (n=30 patients, Table 1, Supplementary Table S1). The effect of mutations and tumor mutation burden in immunotherapy-treated patients was investigated in the MSK-IMPACT study (n=181 patients >59yo (Samstein et al.)).

The exploratory phase of this analysis was conducted using data from the Cancer Genome Atlas (TCGA) (Akbani et al. 2015; Weinstein et al. 2014) and online (https://www.cancer.gov/about-nci/organization/ccg/research/structural-genomics/tcga).

To minimise bias in the data only metastatic melanoma samples were used for this exploration. Clinical and genetic mutation data for all patients with metastases at age >59 or <=55 were included. These experimental age cut-offs were decided *a priori* to obtain more clear-cut young and old age categories; and to ensure that this did not introduce an undue bias we further analysed the data using a simple <59 vs >60 cut-off and a wide interval of <50 vs >70. In both situations the survival in the older age group remained statistically significantly poorer with increasing magnitude as the gap widened. The Stage data in Table 1a in TCGA refers to the stage at diagnosis, but significant numbers of stage 1/2 patients went on to develop metastases in this cohort, and the sequencing data in TCGA was derived from those metastases.

To validate the findings from the exploratory analysis we studied clinical and sequencing data collected as part of rigorous clinical research programs from two separate, unrelated studies in cohort A: La Timone University Hospital, (Aix Marseillle, France) and cohort B: Instituto Valenciano de Oncología (Valencia, Spain) using well established methodology (Gaudy-Marqueste et al. 2014). Relevant institutional review boards and ethics committees at both institutions in both countries approved molecular research and all human participants gave written informed consent. The validation cohorts are unselected, clinical cases prospectively collected without specific targeted gene signatures planned to minimise confirmation bias.

Both cohorts had comprehensive clinical outcome data (supplemental table S1) and both were subjected to targeted deep sequencing at a depth of x100 of 40 melanoma genes from primary melanoma samples. The melanoma panel includes genes found mutated. in >10% of TCGA samples or genes mutated in less than <10% of samples that are known cancer drivers, including *BRAF, NRAS, CDKN2A* and *IDH1*.

Exomes were captured with Agilent SureSelect ALL Exon (V6, V6+UTRs r2) and sequenced on Illumina HiSeq 2000. Sequences were aligned with Burrow-Wheeler Alignment (BWA 0.7.15) with MEM option onto human reference genome, hg19. Base quality was measured with GATK 3.7 tools BaseRecalibrator and PrintReads. Samtools used for variant calling. Somatic mutations were annotated with Oncotator 1.9.0.0. Somatic variants were detected using MuTect2 (GATK 3.7) in the A cohort, and with VarScan 2 in the B cohort. Both sets of variants were filtered to eliminate common germline variants.

The data in Table 1 show the median and range TMB of elderly melanoma samples, which demonstrate overlapping ranges with TCGA samples. Critically, the proportion of the 4-gene driver mutations in the primary tumour B cohort is not statistically different (p=0.09) to the A validation cohort. The 4-gene signature is significantly less frequent in the primary cohorts (p= 0.0008 B cohort, p=0.048 A cohort) than in the metastatic melanoma TCGA cohort, as you would expect for unselected populations of primary melanoma that are not enriched for samples with poor prognosis. However, when compared only in the elderly subgroups the proportions are not significantly different between any of the cohorts (p>0.1 in all comparisons) These three cohorts represent distinct populations of tumours at varying stages of disease. The metastatic tumours in TCGA are from more advanced cancers than the primary cohorts in the validation set, and the 4-gene DNA signature is enriched as these patients represent a poor outcome, metastatic group *per se*. The two primary cohorts are relatively smaller and the heterogeneity of cancer progression and signature within them is likely to be a consequence of this smaller sample size.

#### • Regression modelling

Regression modelling of survival in the metastatic melanoma (TCGA) cohort was conducted using Cox proportional hazards modelling in R implemented in the Survival package. We started with a full model including the following clinical characteristics: age (continuous), sex (binary), patient weight (continuous), Breslow tumour thickness of original primary tumour (continuous) and somatic variant count (log continuous – see below). We initially tested all possible interactions and found no statistically significant interaction effects in this model. To derive the minimum model, we excluded the least statistically significant variable at each step until a minimum model was achieved in which all variables were significant (p<0.05) (supplemental table S2). The variables were excluded in the following order: Weight, Sex, Breslow, variant count. Finally, each excluded variable was added back into the model and checked to ensure that they remained non-significant. To explore the effects of variant count in the model we ran the above sequence initially with variants as a log continuous variable, and then as a binary (high/low) variable with a series of cut-off values (25, 50, 100, 110, 150, 150, 500). Of these only 100 and 110 showed a significant impact in the final model and both were less significant than age in those cases. This finding is consistent with a previous published model from this cohort. All relevant analytic code is available from the authors upon request.

#### • Identification of driver genes

To test the hypothesis that melanoma development in older patients is driven by accumulation of mutations in specific risk genes, we analysed the mutation data using the oncodriveCLUST algorithm (Tamborero et al. 2013) as implemented in the maftools package in R to discover the key driver mutations. This algorithm is a powerful and well-described method validated in the literature and has been used extensively to successfully identify driver mutations/genes in numerous cancer contexts. OncodriveCLUST considers the background mutation rate, gene length and the clustering characteristics of mutations identified in order to derive a confidence value for labelling a gene as a driver as opposed to a passenger. We make no claims regarding function or mechanistic biology behind these mutations but used this bioinformatic tool as a discriminator to identify genes that are likely to be important in the older patient population, but not the younger one. All relevant analytic code is available from the authors upon request.

#### • Gene expression

For the TCGA data, differential gene expression analysis was performed using the DESeq2 (version .28.1) (Love et al. 2014) package in R (version 4.0.0, RStudio v1.2.5003, RStudio Inc). RNA-seq count data for the TCGA SKCM cohort was acquired using the TCGABiolinks package (version 2.16.0 (Colaprico et al. 2015)). The count dataset was filtered to remove any genes that had less than 1000 counts across all samples. Differential expression analysis was run between long term and short-term survivors in the aged (>59 years) metastatic cohort. Long-term survivors were defined as any patient whose overall survival time was 2000 days or greater, and these were compared to patients who died before 2000 days. For pathway enrichment analysis genes that were significantly differentially expressed (FDR p-value < 0.1) were compared against the Reactome Database (Jassal et al. 2020) using the Molecular Signatures Database v7.0 (Msigdb, Broad Institute, https://www.gsea-msigdb.org/gsea/msigdb/index.jsp).

The Lund cohort comprises microarray expression data (Gene Expression Omnibus GSE65904) of 214 melanomas(Cirenajwis et al. 2017), of which 66 are metastatic, with long term survival data. We tested the differential expression analysis between long term and shortterm survivors in the aged cohort as described above. Analysis was performed using the Limma package (version 3.44.1) in R. MCPCounter packaged was used as above to generate cell population scores for each tumor. Where a marker gene had more than one probe quantified on microarray, the most abundantly expressed transcript was used in MCPCounter, that is the probe that had the highest mean value across all samples(Ritchie et al. 2015).

#### • Cell populations

Cell population scores were generated using the MCPCounter R package (version 1.2.0 (Becht et al. 2016)) in the TCGA SKCM RSEM normalised RNA-seq expression data. CD8 T cell scores were compared between long term and short-term elderly metastases using a Mann-Whitney test in GraphPad Prism (version 7.01).

#### • Response to Immunotherapy

To investigate the effects of the driver genes and tumor mutation burden in immunotherapy treated patients data from the MSK-IMPACT study (Samstein et al.) was downloaded from the cBioPortal database (http://www.cbioportal.org/study/summary?id=msk_impact_2017) and reanalyzed. The melanoma cohort included 320 patients, 181 of whom were older than 59 at diagnosis, all of whom had been treated with immunotherapy agents and whose sequencing data included the mutation status of the four driver genes. The sequencing methodology and analysis pipeline for these samples is described in the previous relevant literature (Zehir et al. 2017). To determine high vs low tumor mutation burden in this cohort we investigated the centile groupings within the melanoma group and used the third decile (i.e. those in the highest 30% tumor mutation burden) as high. As did the original MSK-IMPACT authors, we tested a range of deciles (as per supplemental data appendix of the original publication (Samstein et al.)). We tested all deciles and found that 30% was the inflection point at which the response curve separation began to accelerate. Any cutoff from 30% to 10% results in significance (with increasing magnitude as the cutoff becomes more stringent) – we chose a cutoff at 30%, that included as many patients as possible.

#### Data and materials availability

All data used are either public, or (for patient-level data) available without restriction in anonymized form from the authors. The analytical code used is available from the authors without restriction except to anonymize patient identifiable information.

#### Supplementary Figures

**Figure S1.**
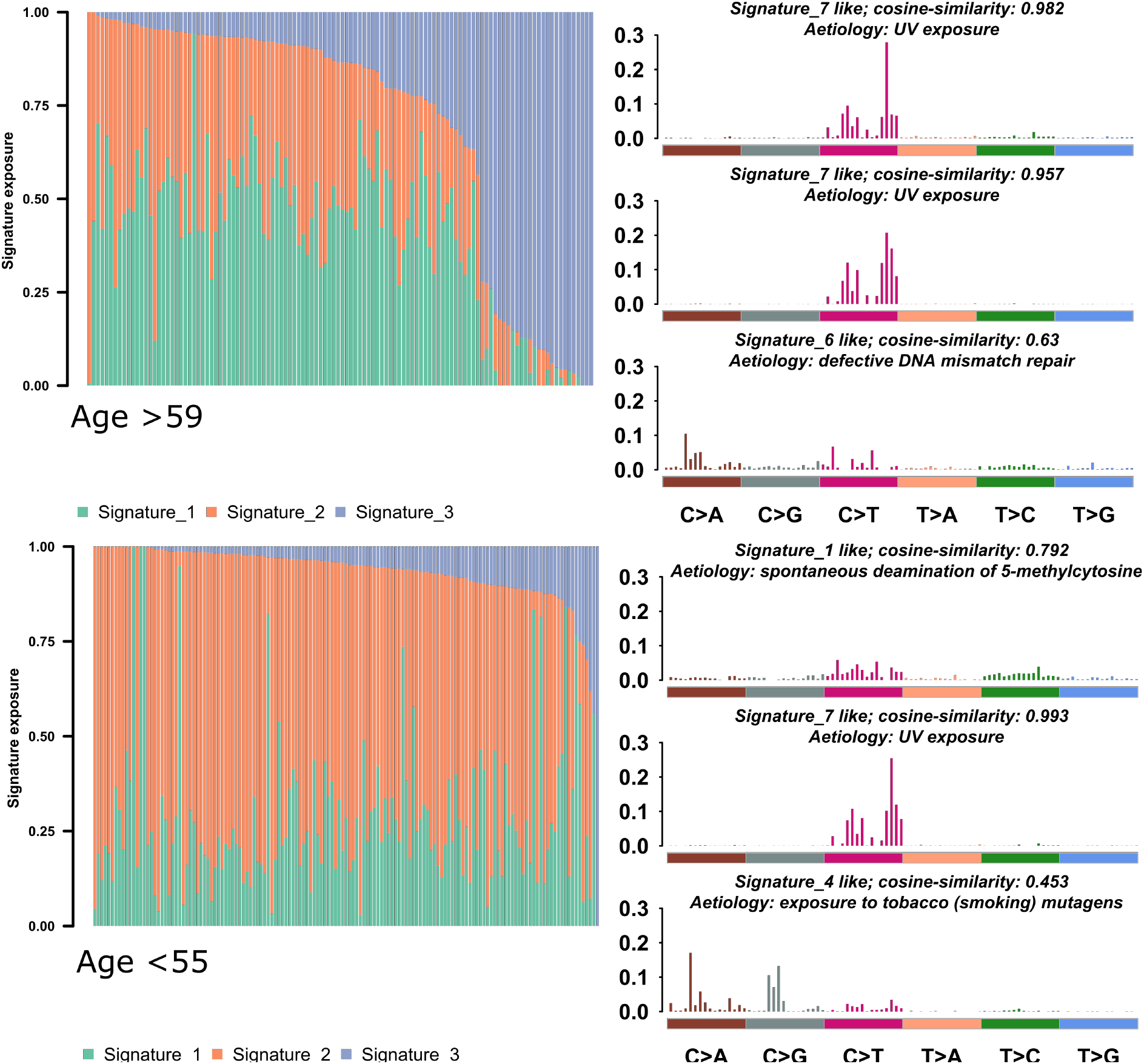
Mutational signature decomposition old and young melanoma samples from TCGA. The vast majority of the mutational contribution comes from UV damage, with minor contributions that differ between the two groups.

**Figure S2.**
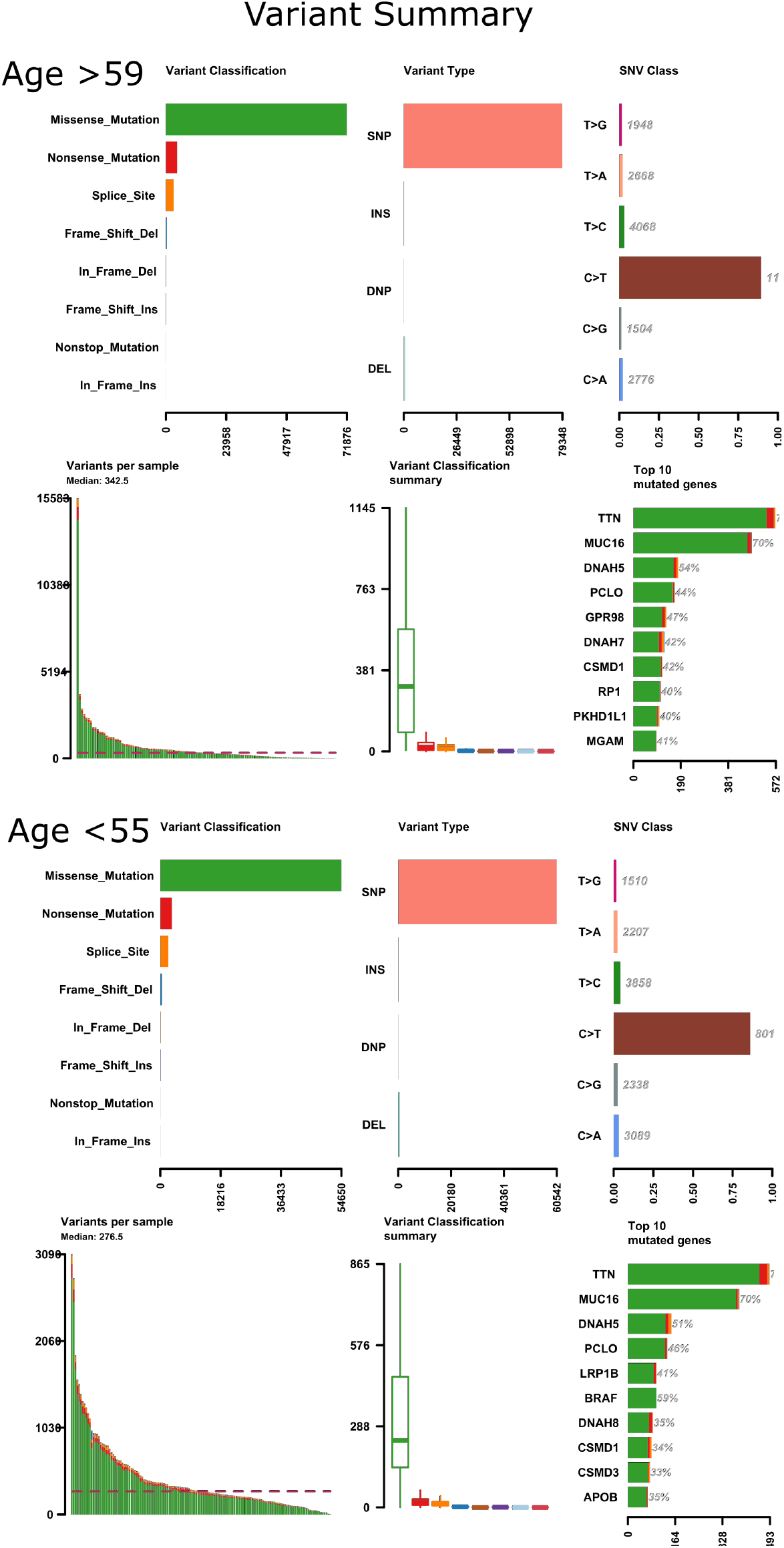
Summary of the types and frequencies of mutations in metastatic melanoma samples from old or young patients in TCGA. Older patients harbour greater numbers of mutations in all categories, but the relative proportions of types of mutations are similar between the groups.

**Figure S3.**
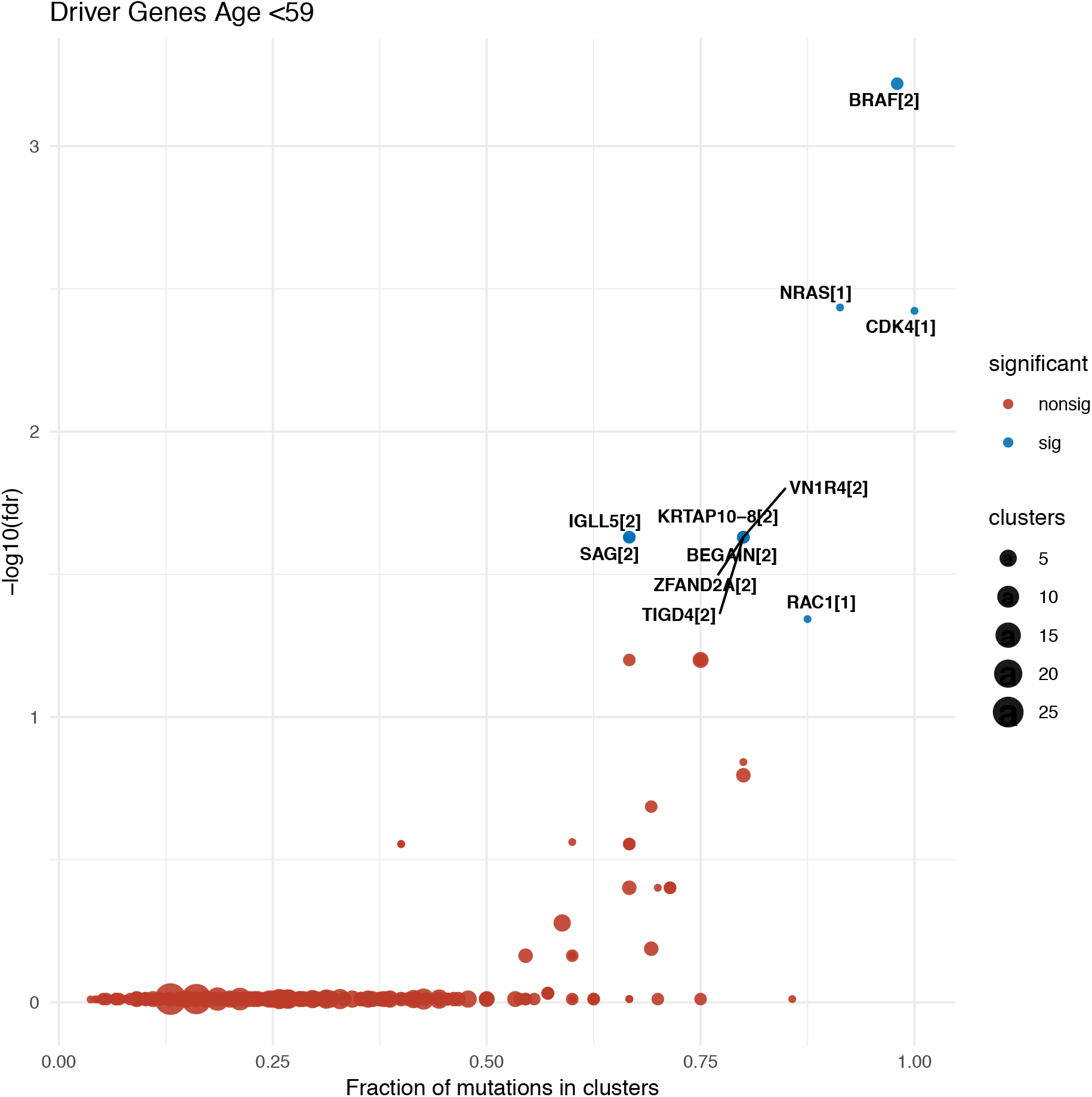
OncodriveCLUST analysis of metastatic melanoma in age <55 showing 11 driver genes and their clusters in brackets.

**Figure S4.**
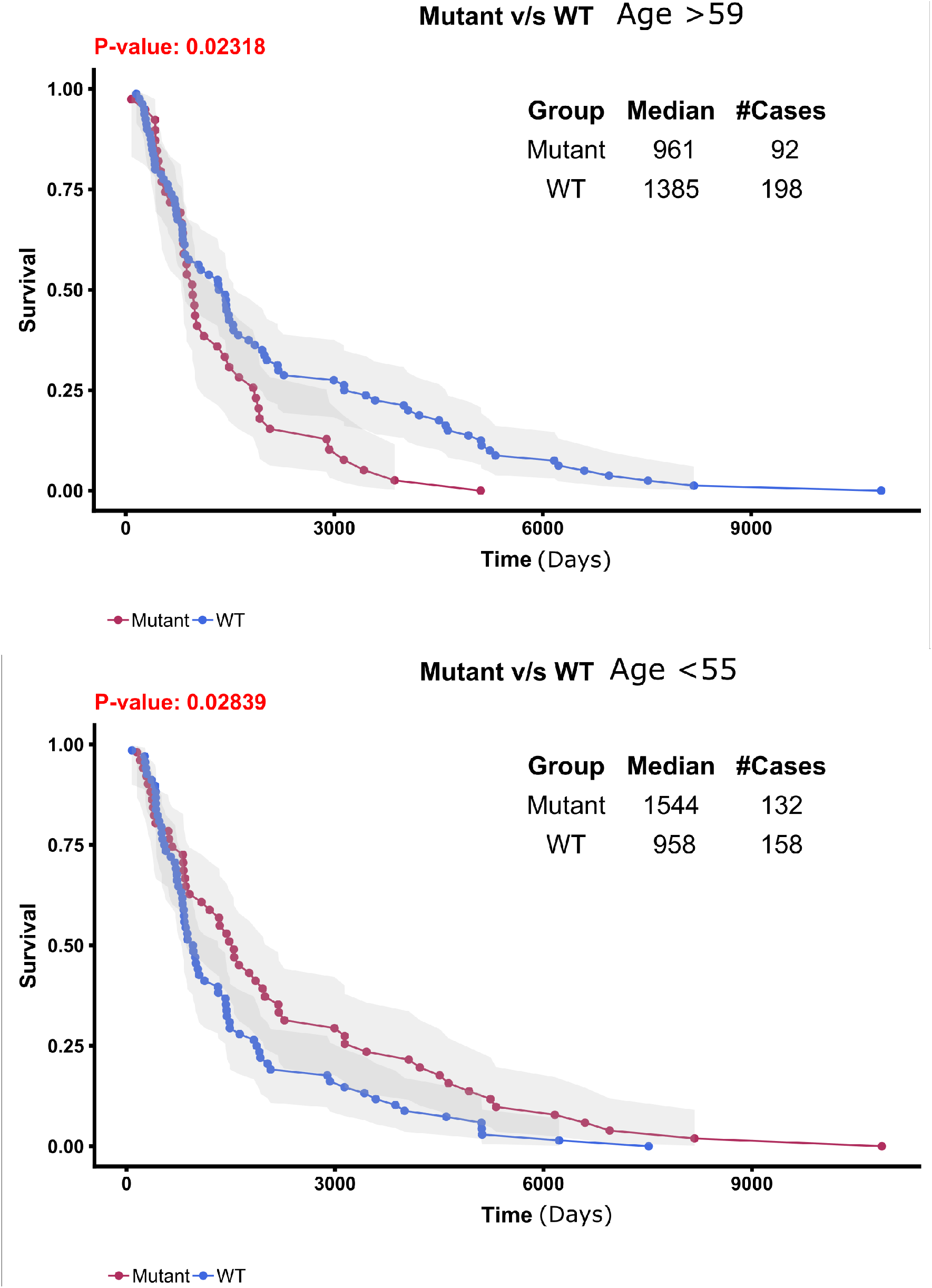
Overall survival in metastatic melanoma in TCGA. Upper plot shows the predictive effect of mutations in any of the 18 identified driver mutations in the elderly, the lower plot show the same for the 11 driver mutations in young melanoma.

**Figure S5.**
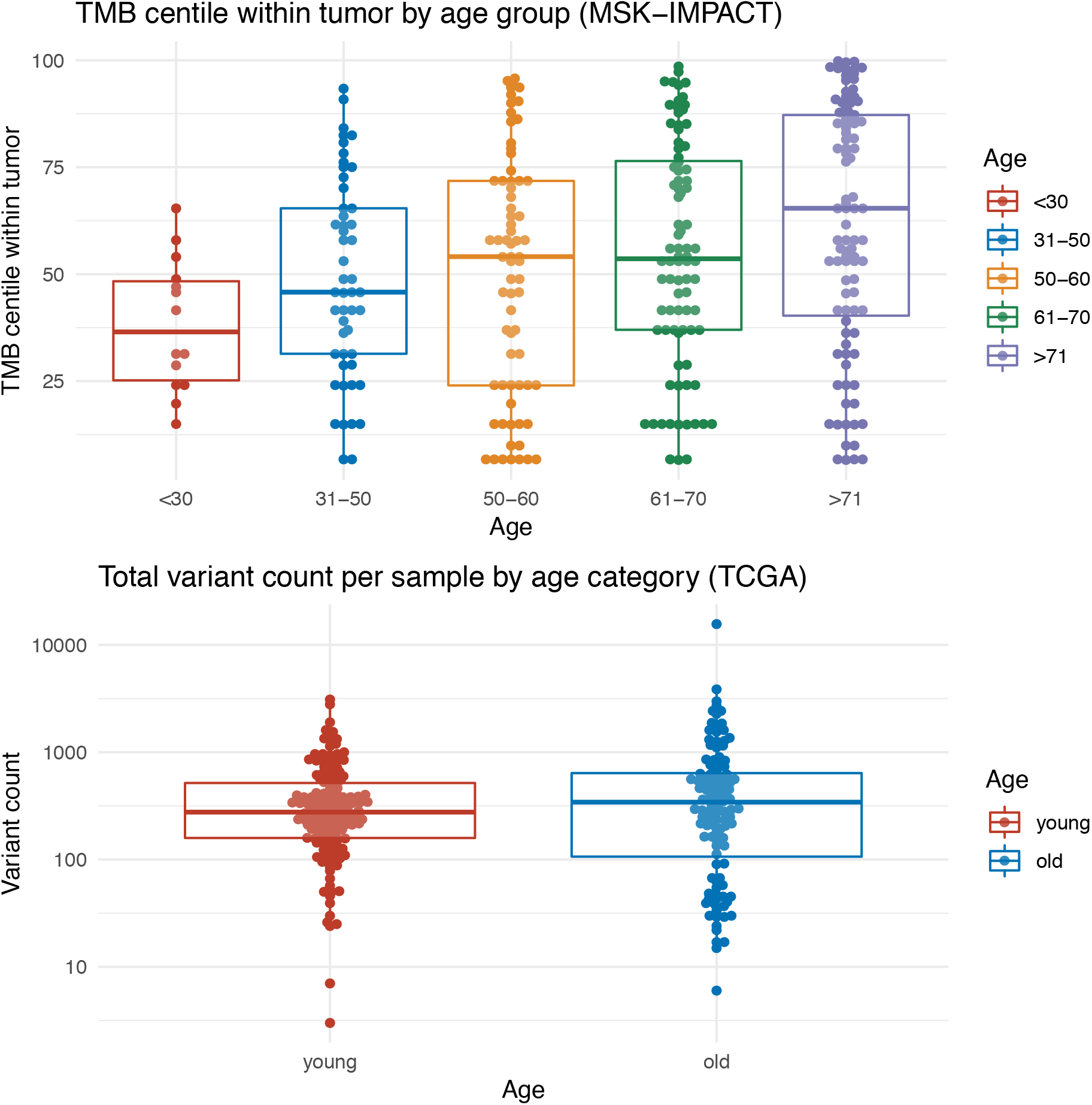
Correlation between age and TMB across in the TCGA and MSK-IMPACT cohorts.

#### Supplementary tables

**Table S1.**
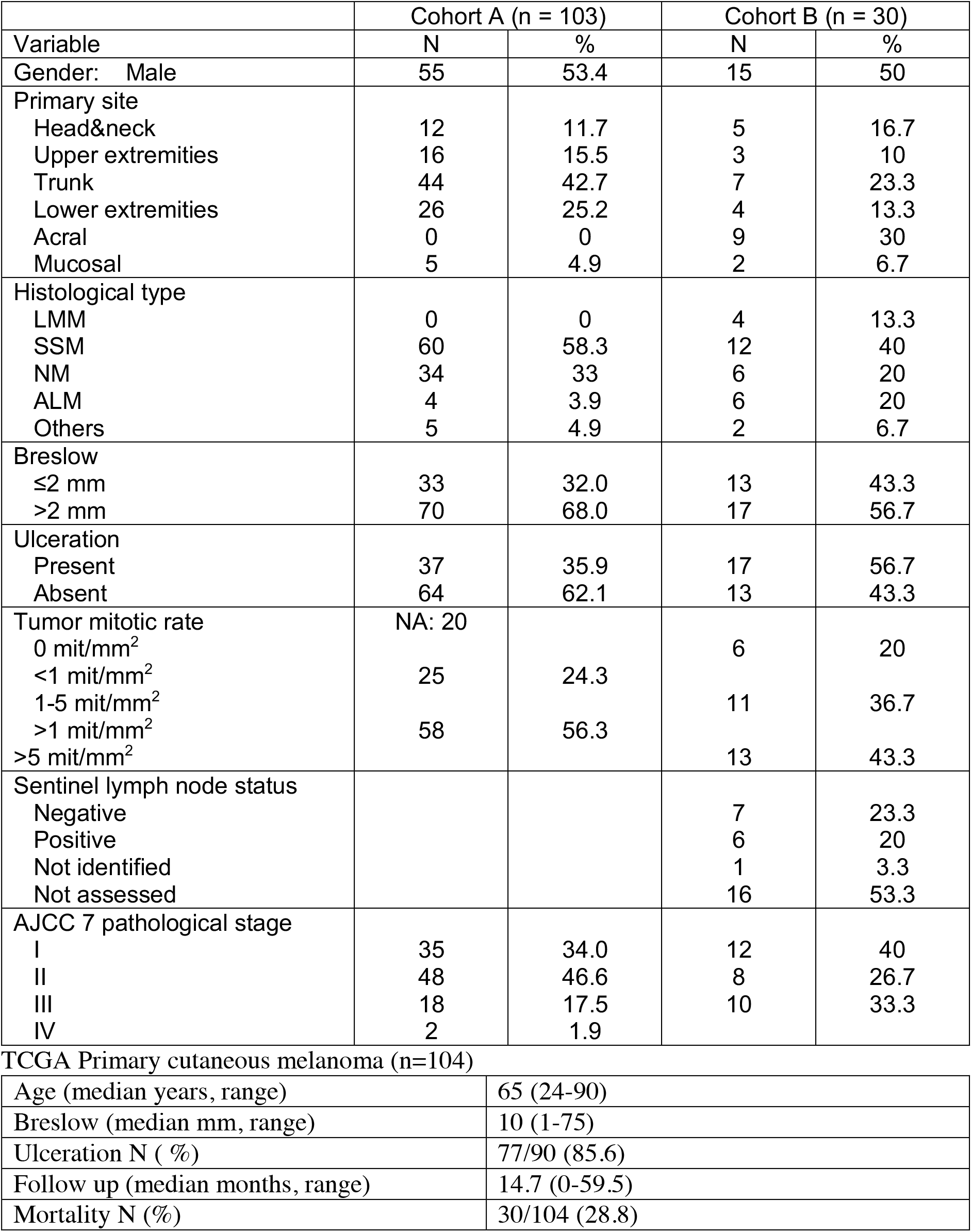
Full clinical details of the patients included in the analysis of the elderly (age >= 60) A and B primary melanoma cohorts and TCGA primary melanoma cohort.

|  | Cohort A (n = 103) |  | Cohort B (n = 30) |  |
| --- | --- | --- | --- | --- |
| Variable | N | % | N | % |
| Gender: Male | 55 | 53.4 | 15 | 50 |
| Primary site |  |  |  |  |
| Head&neck | 12 | 11.7 | 5 | 16.7 |
| Upper extremities | 16 | 15.5 | 3 | 10 |
| Trunk | 44 | 42.7 | 7 | 23.3 |
| Lower extremities | 26 | 25.2 | 4 | 13.3 |
| Acral | 0 | 0 | 9 | 30 |
| Mucosal | 5 | 4.9 | 2 | 6.7 |
| Histological type |  |  |  |  |
| LMM | 0 | 0 | 4 | 13.3 |
| SSM | 60 | 58.3 | 12 | 40 |
| NM | 34 | 33 | 6 | 20 |
| ALM | 4 | 3.9 | 6 | 20 |
| Others | 5 | 4.9 | 2 | 6.7 |
| Breslow |  |  |  |  |
| ≤2 mm | 33 | 32.0 | 13 | 43.3 |
| >2 mm | 70 | 68.0 | 17 | 56.7 |
| Ulceration |  |  |  |  |
| Present | 37 | 35.9 | 17 | 56.7 |
| Absent | 64 | 62.1 | 13 | 43.3 |
| Tumor mitotic rate | NA: 20 |  |  |  |
| 0 mit/mm<sup>2</sup> |  |  | 6 | 20 |
| <1 mit/mm<sup>2</sup> | 25 | 24.3 |  |  |
| 1-5 mit/mm<sup>2</sup> |  |  | 11 | 36.7 |
| >1 mit/mm<sup>2</sup> | 58 | 56.3 |  |  |
| >5 mit/mm<sup>2</sup> |  |  | 13 | 43.3 |
| Sentinel lymph node status |  |  |  |  |
| Negative |  |  | 7 | 23.3 |
| Positive |  |  | 6 | 20 |
| Not identified |  |  | 1 | 3.3 |
| Not assessed |  |  | 16 | 53.3 |
| AJCC 7 pathological stage |  |  |  |  |
| I | 35 | 34.0 | 12 | 40 |
| II | 48 | 46.6 | 8 | 26.7 |
| III | 18 | 17.5 | 10 | 33.3 |
| IV | 2 | 1.9 |  |  |

**Table S1.** Full clinical details of the patients included in the analysis of the elderly (age ≥ 60) A and B primary melanoma cohorts and TCGA primary melanoma cohort.
|  |  |
| --- | --- |
| Age (median years, range) | 65 (24-90) |
| Breslow (median mm, range) | 10 (1-75) |
| Ulceration N ( %) | 77/90 (85.6) |
| Follow up (median months, range) | 14.7 (0-59.5) |
| Mortality N (%) | 30/104 (28.8) |

**Table S2.** Multivariate regression using a cox proportional hazards model. Non-significant variables sequentially eliminated from the model during iterative process

|  | Iteration 1<br>(n=184, events: 85) |  | Iteration 2<br>(n=184, events: 85) |  | Iteration 3<br>(n=231, events: 103) |  | Iteration 4<br>(n=287, events: 125) |  |
| --- | --- | --- | --- | --- | --- | --- | --- | --- |
|  | Hazard<br>ratio | p-value | Hazard<br>ratio | p-value | Hazard<br>ratio | p-value | Hazard<br>ratio | p-value |
| Age <55 | 0.49 | 0.004 | 0.49 | 0.004 | 0.50 | 0.001 | 0.52 | 0.001 |
| AJCC IV | 3700 | 1.0 | 3700 | 1.0 | 5.1 | 0.027 | 4.79 | 0.032 |
| Log (TMB) | 0.99 | 0.40 | 0.99 | 0.40 | 1.0 | 0.48 |  |  |
| Breslow | 1.0 | 0.06 | 1.0 | 0.06 |  |  |  |  |
| Male sex | 1.0 | 0.96 |  |  |  |  |  |  |

